# Conformational flexibility of GRASP protein and its constituent PDZ subdomains reveals structural basis of its promiscuous interactome

**DOI:** 10.1101/666495

**Authors:** Luis Felipe S. Mendes, Mariana R. B. Batista, Peter J. Judge, Anthony Watts, Christina Redfield, Antonio J. Costa-Filho

## Abstract

The Golgi complex is a central component of the secretory pathway, responsible for several critical cellular functions in eukaryotes. The complex is organized by the Golgi matrix, which includes the Golgi Reassembly and Stacking Proteins (GRASPs), which participate in cisternae stacking and lateral linkage in vertebrates. GRASPs also have critical roles in other processes, with an unusual ability to interact with several different protein binding partners. The conserved N-terminus of the GRASP family includes two PDZ domains. Previous crystallographic studies of orthologues suggest that PDZ1 and PDZ2 have similar conformations and secondary structure content, however PDZ1 alone mediates nearly all the interactions between GRASPs and their binding partners. In this work, NMR, Synchrotron-Radiation Circular Dichroism and Molecular Dynamics were used to examine the structure, flexibility and stability of the two constituent PDZ domains. GRASP PDZs are structured in an unusual β_3_α_1_β_4_β_5_α_2_β_6_β_1_β_2_ secondary structural arrangement and NMR data indicates that the PDZ1 binding pocket is formed by a stable β_2_-strand and a more flexible and unstable α_2_-helix, suggesting an explanation for the higher PDZ1 promiscuity. The conformational free energy profiles of the two PDZ domains were calculated using Molecular Dynamics simulations. The data suggest that, after binding, the protein partner significantly reduces the conformational space that GRASPs can access by stabilizing one particular conformation, in a partner-dependent fashion. The structural flexibility of PDZ1, modulated by PDZ2, and the coupled, coordinated movement between the two PDZs enable GRASPs to interact with multiple partners, allowing them to function as promiscuous, multitasking proteins.

**Significance Statement:** Golgi Reassembly and Stacking Proteins (GRASPs) play pivotal roles in the maintenance of Golgi structure as well as in unconventional protein secretion. Their broad network of interactions is mainly sustained by the two-PDZ domains located in the N-terminal portion of the protein. The asymmetry of the PDZ domains in terms of number and diversity of interacting partners has been long recognized, but the molecular determinants of that asymmetry remains largely unknown. The biophysical data presented here provide a firm basis for understanding why PDZ1 behaves differently to PDZ2 in solution, despite their similar 3D structures. Furthermore, we propose that PDZ2 assist ligand binding to PDZ1, by means of conformational stabilization.

## Introduction

The Golgi complex is a membranous organelle located in the heart of the secretory pathway, responsible for several critical cellular functions [1,2]. In vertebrates, this organelle is assembled as a stack of flattened cisternae that can also be laterally connected in a ribbon-like shape [1]. Visually, the Golgi complex is one of the most immediately recognizable organelles inside higher eukaryotic cells. The complex is held together by proteins that constitute the so-called Golgi matrix, however the mechanism by which they accomplish this task is not fully understood [3]. It is likely that proteins belonging to the Golgi matrix were present in the last eukaryotic common ancestor (LECA), suggesting that eukaryotic cells share some of the mechanisms controlling the Golgi structure [4]. One of the most important proteins of the Golgi matrix (probably present in LECA [4]), is the family of peripheral membrane proteins called Golgi Reassembly and Stacking Proteins (GRASPs) [5].

GRASPs were first observed to be essential factors for Golgi reassembly *in vitro* [6,7] and *in vivo* [8]. They participate in the lateral linkage of Golgi stacks in vertebrates (building the Golgi ribbon), and also in all the Golgi remodelling necessary in migrating cells [9]. Current models suggest that GRASP-mediated Golgi stacking occurs via GRASP dimerization in a *trans* orientation [10], whereby a monomeric GRASP on one cistern face interacts with a second GRASP monomer on an opposite cistern [10]. The mechanism of GRASP dimerization (and whether higher-order oligomers exist and might play any role) is still unclear and of intense debate [10,11,12].

Although first identified as stacking factors, other functions of GRASPs have also been observed. GRASP can interact directly with the C-terminus of the Transforming Growth Factor-α (TGF-α) and with several members of the p24 family, facilitating the conventional protein secretion pathway [13,14]. GRASPs also have a direct and essential role in the unconventional secretion of a large number of proteins, such as: (1) the soluble acyl-coenzyme A binding protein (ACBP) in *Dictyostelium* [15], *Saccharomyces cerevisiae* and *Pichia pastoris* [16,17], (2) the Golgi bypass of αPS1 integrin during the stage 10B of *Drosophila* embryogenesis [18], (3) the mutant ΔF508 of the cystic fibrosis transmembrane conductance regulator (CFTR) in an ER stress situation [19], (4) in the secretion of IL1-β [20], among others [21,22]. GRASP is apparently also required for the export of non-protein molecules. In *Cryptococcus neoformans*, deletion of GRASP (CnGRASP) results in inefficient secretion of its main virulence factor GXM, followed by reduced capsule formation and attenuated virulence [23].

Although many functions associated with GRASPs have been identified, their detailed molecular mechanisms are still unclear. The crystal structures of the N-terminal domain of human GRASP55 (24), *Rattus norvegicus* GRASP55 (25) and *Rattus norvegicus* GRASP65 (25) have only recently been reported. GRASPs are composed of two structurally similar PDZ domains (named after the three first PDZ-proteins found: <u>P</u>SD-95, <u>D</u>LG and <u>Z</u>O-1) with an atypical protozoa-like arrangement, which together comprise the so-called GRASP domain (DGRASP). This finding also suggests that a new sequence analysis criterion might reveal a host of unidentified eukaryotic PDZ domains [24]. A second region in the GRASP structure consists of a poorly conserved domain that is rich in serine and proline (SPR domain) and that has a regulatory function [26,27] and intrinsically disordered characteristics [28].

One of the most intriguing properties of GRASPs is their ability to interact with a large variety of different protein partners. In addition to the Golgins, mammalian GRASPs also interact with cargo receptors from the conventional secretory pathway [13] and form part of a complex with RAB2, essential for normal protein transport and Golgi structure [29]. GRASPs have also been shown to interact with TGF-α [14], the potassium channel tetramerization domain-containing protein 5 (KCTD5) [30], with SEC16A and ΔF508 CFTR in response to ER stress [31,19], with JAM-B and JAM-C during spermatogenesis [32], with LC3-II on the autophagosomes and LAMP2 on late endosomes/lysosomes [33], and also with itself during dimerization [34]. These interactions involve the PDZ1 domain rather than PDZ2, despite the structural similarity seen in the reported crystal structures [11,12,19,25,32]. PDZ1 was also observed in crystal structures to be sufficiently promiscuous to interact with a non-native truncated SPR C-terminus [25].

In a previous report, we showed that GRASPs from the fungal pathogen *C. neoformans* [28] and *S. cerevisiae* [35] present structural features usually observed in molten globule proteins, even in the absence of any mild denaturing conditions. As expected for a collapsed intrinsically disordered protein, many physicochemical perturbations induced by the crystal packing, (including local dehydration and changes in the dielectric constant, along with increased PEG concentration used in the crystallographic reservoir), promote disorder-to-order transitions in GRASPs [35,36]. Even though GRASPs appear to have a special affinity for di-valine-containing C-terminal proteins [27], the overall sequences of their protein partners do not have a conserved sequence or structural feature in common (Figure S1), suggesting GRASPs might be highly promiscuous in terms of protein-protein interactions. Therefore, this apparent functional asymmetry of the PDZs and their contrasting promiscuity of interactions with binding partners, might benefit from detailed studies of GRASPs in solution.

Since crystal structures have not provided a clear rationale as to why the two PDZ subdomains behave so differently *in vivo*, we used high-field solution-state nuclear magnetic resonance (NMR), synchrotron radiation circular dichroism (SRCD), conventional circular dichroism (CD), steady-state fluorescence and molecular dynamics (MD) simulations to probe the flexibility and stability of different parts of the GRASP structure. Our data show that the two PDZs behave differently in solution, with most of the flexible and promiscuous regions located right in the binding pocket of PDZ1, whereas PDZ2 is significantly more ordered. The coordinated movement between the two subdomains covers a wide conformational space, accessible from the free energy present at physiological temperatures, which collapses to a small area after binding with the protein partner. We propose that the orchestrated movement of the PDZs is used by the protein to control access to the PDZ1 binding pocket, and an array of secondary interactions with PDZ2 dictates the overall stability and the strength of the protein-ligand complex. Our data give a convincing explanation as to why PDZ1 is capable of mediating almost all the interactions with other protein partners, and suggest a new role for GRASPs as highly promiscuous, multitasking proteins.

## Materials and Methods

### Protein expression and purification

CnGRASP GRASP domain (DGRASP) and isolated PDZ samples were expressed and purified as described elsewhere [28]. ^13^C- and/or ^15^N-labeled DGRASP and PDZs were produced using a protocol reported previously [37]. The same protocol was used for purification of both labelled and unlabelled protein [28].

### Circular Dichroism (CD)

CD measurements were performed using a Jasco J-815 Spectropolarimeter fitted with a Peltier temperature control unit. High-grade quartz cuvettes of 1 mm path length were used for data collection. The temperature was 20 °C. Further parameters were: data pitch 0.5 nm, D.I.T. of 1 sec, 1.00 nm bandwidth and a scanning speed of 50 nm/min. The sample preparation for the urea titration consisted of a protein dilution (less than 5% of the total final volume) directly into a solution containing 50 mM sodium phosphate, 10 mM NaCl and 1 mM β-mercaptoethanol, pH 7.4 with the corresponding urea concentration. Protein concentration was kept constant at 0.15 mg/mL.

### Synchrotron Radiation Circular Dichroism (SRCD)

SRCD experiments were performed on the B23 Synchrotron Radiation CD beamline at Diamond Light Source, Oxfordshire, UK. Protein concentration was in the range of 5-10 mg/mL and demountable quartz cells of 20 µm and 50 µm were mainly used. The parameters were the same as those used for the conventional CD experiments.

### Fluorescence Spectroscopy

Steady-state fluorescence was monitored using a Hitachi F-7000 spectrofluorimeter equipped with a 150 W xenon arc lamp and polarized filters for anisotropy experiments. The excitation and emission monochromators were set at 5 nm slit width in all tryptophan experiments and 5 nm excitation with 10 nm emission for the 1-anilino-8-naphthalenesulfonic acid (ANS) experiments. For tryptophan fluorescence experiments, the selective tryptophan excitation wavelength was set at 295 nm and the emission spectra were measured from 310 up to 450 nm. For the anisotropy measurements, tryptophan was selectively excited at 300 nm and the mean anisotropy values were calculated as the mean average of the values from ±10 nm (1 nm step acquisition) over the wavelength of maximum emission determined for each condition. For the ANS (250 µM) fluorescence experiments, the excitation wavelength was set at 360 nm and the emission spectra were measured from 400 up to 650 nm. The urea unfolding experiments were performed using the same strategy described in the CD experiments. The first moment of the fluorescence spectrum, or spectral center of mass, is calculated according to the equation 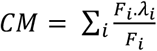 where *F*_*i*_ is the fluorescence intensity of the particular wavelength *λ*_*i*_. All the experiments were performed at 25°C.

### High-Field Solution Nuclear Magnetic Resonance (NMR)

All NMR experiments were carried out using spectrometers operating at ^1^H frequencies of 600 MHz and 950 MHz, equipped with Oxford Instruments magnets and Bruker Avance III HD consoles. The 600 MHz and 950 MHz spectrometers were equipped with a 5mm TCI cryoprobe and a 5 mm room-temperature TXI probe, respectively. Data were processed using NMRPipe [38] and/or Bruker TopSpin 3.5 and spectra were analyzed using the CCPN software [39]. ^1^H-^15^N HSQC spectra of DGRASP in 25 mM HEPES, 100 mM NaCl, 5 mM β-mercaptoethanol, pH 7.0 (95% H_2_O/5% D_2_O) and in a urea concentration gradient ranging from 0 to 9 M (1 M steps) were collected. The sample with 9 M urea was the one with the highest number of resonance peaks in the ^1^H-^15^N HSQC spectrum and, therefore, chosen for assignment. CBCACONH, CBCANH, HNCO, HN(CA)CO, (H)CC(CO)NH, HBHA(CBCACO)NH, HSQC-TOCSY, HSQC-NOESY, HSQC-NOESY-HSQC were collected for assignment; these were transferred back to all other HSQC spectra collected with lower urea concentrations. The temperature was 20 °C in all experiments and the protein concentration varied from 150 to 400 µM. NMR data collection for PDZ1 and PDZ2 followed the same protocols and buffers used with the DGRASP construction. Shigemi tubes were used in all experiments. The assignments have been deposited in the BioMagResBank (http://www.bmrb.wisc.edu) under the accession number 27921.

### Molecular Dynamics (MD) simulations

The crystal structure of DGRASP55 in complex with the JAM-C C-terminus peptide (PDB ID: 5GMI) was used to build models for holo-DGRASP55 (in which the peptide was present) and apo-DGRASP55 (from which the peptide was absent). The structures were solvated with Packmol [40,41] using ca. 19,500 water molecules and Na^+^ and Cl^−^ ions were added to render the system neutral. Simulations were performed with NAMD [42] using periodic boundary conditions, the CHARMM36 [43] force-field for the protein and peptides, and the TIP3P model [44] for water. Equilibration and simulations were performed in the NPT ensemble at 298.15 K and 1 atm. Temperature was controlled using Langevin dynamics with a friction coefficient of 10 ps^−1^. The Nosé-Hoover algorithm was used for the pressure control, with a piston oscillation period of 200 fs and decay rate of 100 fs. A 2 fs time-step was used to integrate the equations of motion using the Verlet algorithm. A cutoff of 12 Å was used for Van der Waals interactions, whereas the long-range electrostatic interactions were handled by the Particle Mesh Ewald method [45]. Covalent bonds involving hydrogen atoms in the protein were constrained to their equilibrium distances using the SHAKE algorithm, while SETTLE [46] was used for water.

The systems were equilibrated as follows: 1) 1,000 steps of conjugate-gradient steps (CG) [47] followed by 200 ps MD keeping all protein and peptide atoms fixed, for solvent relaxation. 2) 500 CG minimization steps followed by 200 ps MD with fixed Cα coordinates, allowing the side-chains of the protein to relax. 3) 2 ns MD without any restrictions. Five unbiased simulations with 50 ns were performed for each system. The final structures generated, were used for unbiased simulations and for Adaptive Biasing Force (ABF) simulations.

The ABF [48,49] simulations were performed using NAMD. The reaction coordinates were defined as the distance between the center of mass of the two PDZs and the dihedral angle formed by the center of mass of the PDZs and the helix, connecting the two domains (Figure S2). The first reaction coordinate was sampled by ABF simulations within 20 and 35 Å, with a precision of 0.1 Å, and the second reaction coordinate was sampled within −120° and 50°. To restrain the movements of the atoms inside the regions of interest, a harmonic boundary potential with 10 kcal.mol^−1^Å^2^ force constant was applied. The ABF force was applied only after 5000 samples of the mean force were generated in each bin. Ten independent ABF simulations with 100 ns each were performed for each system. The convergence of the free energy profiles was checked by computing the root mean square of the ABF forces. The profiles were considered converged if their root mean square was smaller than 0.2 kcal/mol. VMD [50] was used for visualization and figure preparation.

## Results and Discussion

### DGRASP as a molten globule-like protein

^15^N-or ^15^N/^13^C-labeled and unlabeled samples were prepared using the protocol described in the Materials and Methods section and protein purity was monitored using SDS-PAGE (Figure 1A). CnGRASP, like the orthologue GRASP from *S. cerevisiae* [35], has previously been identified as a molten globule-like protein in solution, and belongs to the family of collapsed intrinsically disordered proteins [28]. We have previously suggested that the isolated DGRASP domain of CnGRASP shares some of the molten globule features observed in the full-length structure [28]. Here we extended that analysis by showing that molten globule-like behavior is indeed observed for the isolated GRASP domain (DGRASP) (Figure 1). In the absence of urea, native DGRASP bound the fluorophore ANS with high affinity, compared to the fully unfolded protein (Figure 1B), consistent with the partitioning of ANS molecules into the hydrophobic interior in the native state of the protein. The gradual changes in the ANS fluorescence spectra at different urea concentrations indicate that the hydrophobic interior unfolds in a linear manner, exhibiting low cooperativity (Figure 1C). This low cooperativity during unfolding was also observed in the loss of regular secondary structure, as monitored by CD (Figure 1D) and in the change of the tryptophan fluorescence anisotropy (Figure 1E). Our data indicate that DGRASP is a molten globule-like protein, and that its three-dimensional structure unfolds in a non-cooperative way.

**Figure 1:**
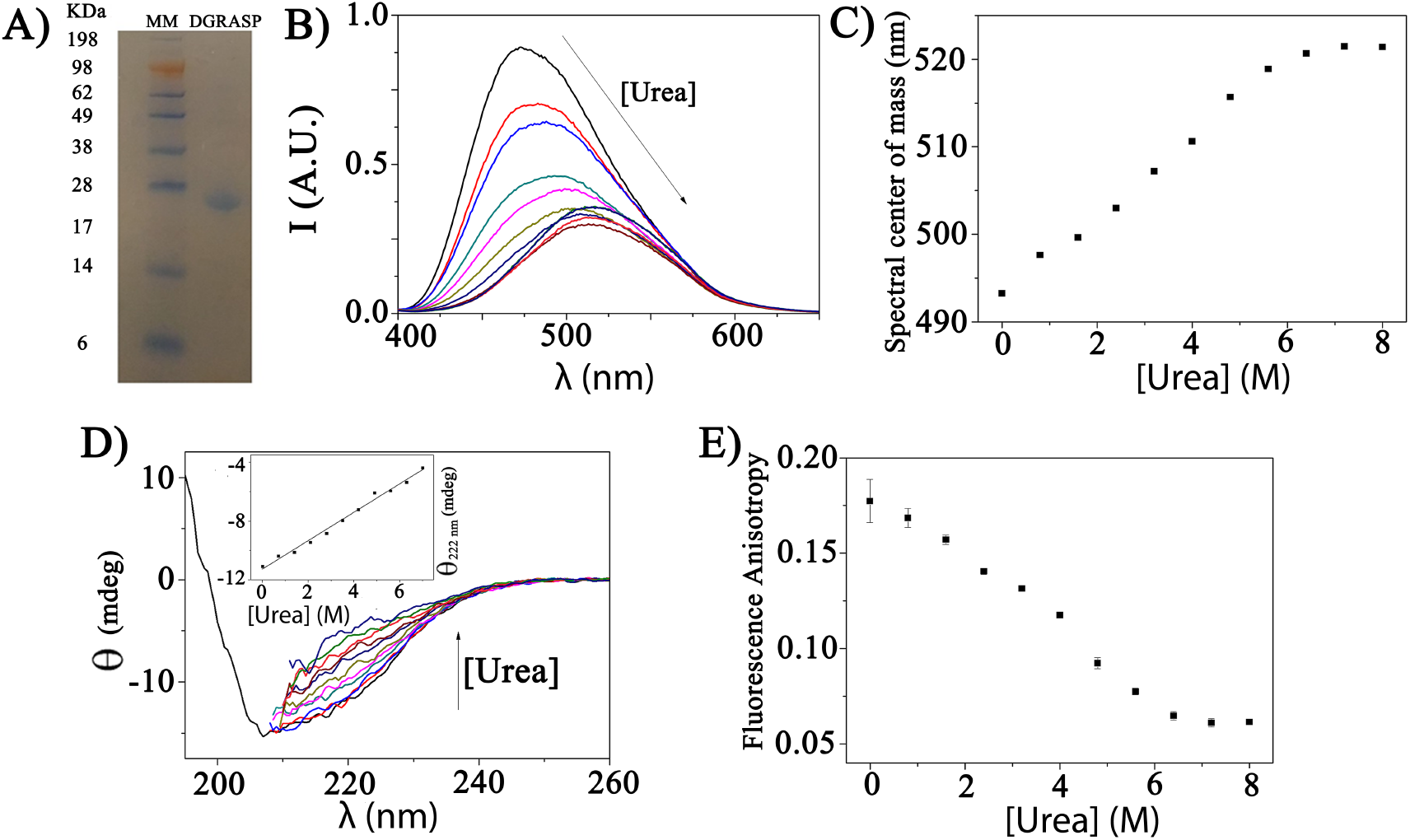
DGRASP as a molten globule-like protein: A) SDS-PAGE of purified ^15^N/^13^C-labeled DGRASP. Lane 1: molecular mass standards in kDa. Lane 2: purified DGRASP. B) ANS-based fluorescence. The arrow indicates the increasing urea concentration. C) The center of spectral mass is calculated from the ANS-based fluorescence spectra and plotted as a function of the urea concentration. D) DGRASP urea titration monitored by CD. Far-UV CD curves for each step of the titration (0.7 M urea from 0 to 7 M). The inset shows the CD intensity at 222 nm plotted as a function of the urea concentration. The dataset was fitted to a linear function. E) Variations in fluorescence anisotropy as a function of urea concentration.

Although intrinsically disordered proteins (IDPs) may have significant secondary structure content, their hydrophobic cores are not stable. IDPs therefore have a three-dimensional conformation that fluctuates over a timescale of µs-ms [51,52]. IDPs are frequently observed as important components of the cellular signaling machinery [53] and may function as central hubs [54], especially due to their unique structural plasticity [51]. It is becoming clear that the understanding of the structural behavior of molten globules in solution is of pharmacological importance. For example, prion-forming sequences, which are especially enriched in asparagine, have been shown to promote molten globule-like structures, in which amyloid-nucleating contacts can be made [55,56]. Mutant forms of the tumor suppressor protein, p53, have recently been shown to behave as molten globules while exhibiting prion-like properties and, although its standard functions are strongly connected to tumor suppression, p53 mutants and aggregates are involved in cancer progression [57,58].

### PDZ asymmetry

It has been previously observed that PDZs in tandem might assist each other during folding [59,60]. For example, for the PDZ12 tandem of the GRIP1 PDZ1–3 cassette, the folding of PDZ1 strictly depends on the covalent attachment to PDZ2 [59]. In addition, PDZ5 of GRIP is completely unstructured in solution but, when covalently bound to PDZ4, both PDZ domains become well folded and stable [60]. We explored the behavior of the PDZs of DGRASP in solution using NMR methods. The DGRASP ^1^H-^15^N HSQC spectrum showed around 10% of the total expected resonance lines, as previously observed for other induced molten globule-proteins (Figure 2 – 0 M urea) [52,61]. Furthermore, the DGRASP ^1^H-^15^N HSQC spectrum showed low proton dispersion (separation of less than 0.7 ppm), a strong indication that those peaks might be located in disordered, highly dynamic regions.

**Figure 2:**
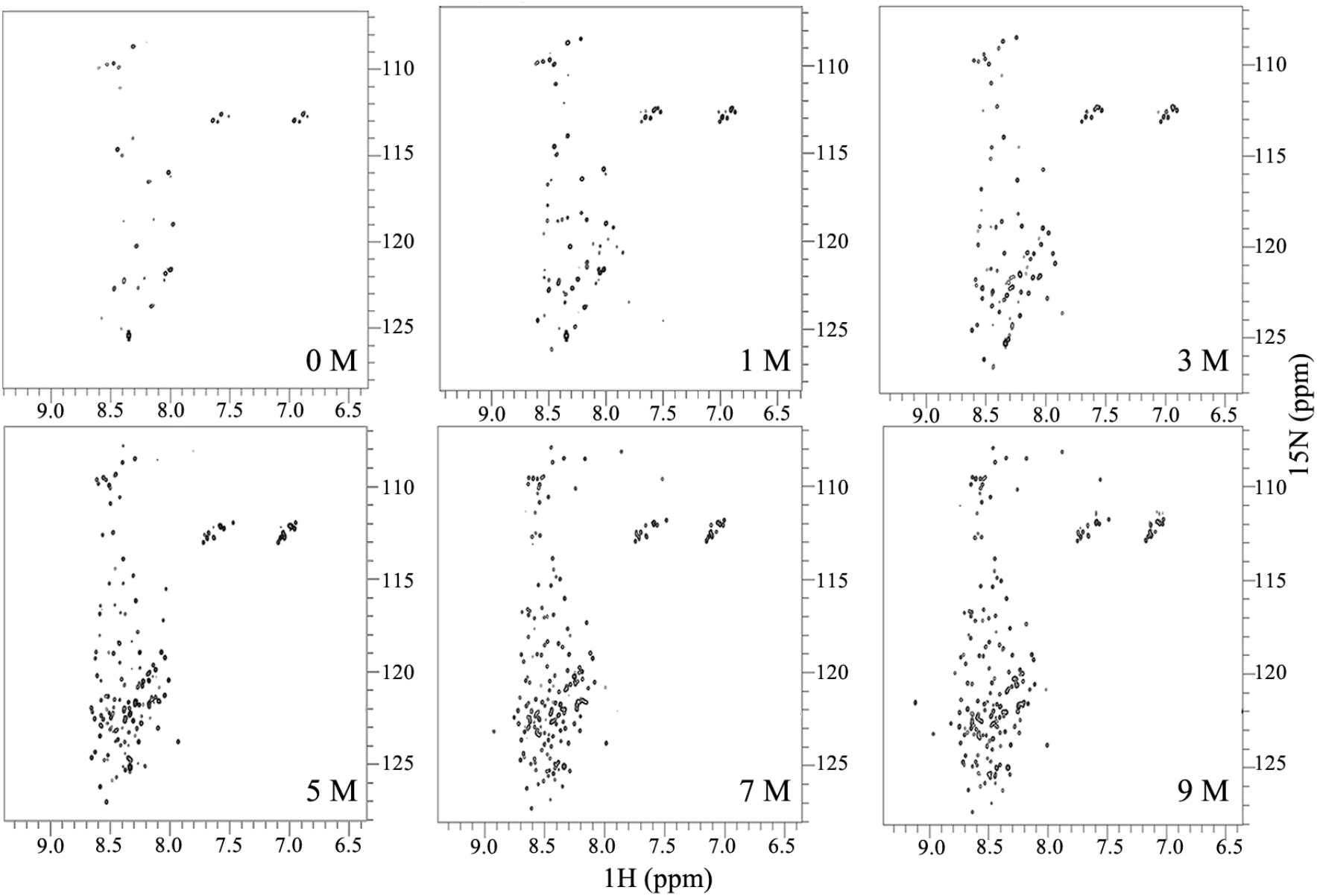
DGRASP urea titration monitored by ^1^H-^15^N HSQC at 950 MHz at 20 °C. The sample was titrated from 0 to 9 M in 1 M steps. Representative spectra are presented.

The absence of resonance lines in the ^1^H-^15^N HSQC spectrum of DGRASP, is consistent with the presence of a molten globule-like fluctuating structure in the characteristic timescale window of µs-ms [52]. To recover the missing peaks, the structure is disrupted by increasing urea concentration, to either increase the local dynamics or decrease the number of “conformational states” the protein visits. The strategy of using urea unfolding to gain structural information by NMR has been previously used to study the molten globule states of α-lactalbumin and p53 [62,63].

Urea was titrated from 0 to 9 M in 1 M steps and selected ^1^H-^15^N HSQC spectra are shown in Figure 2. The resonance peaks start to appear during the titration indicating that the corresponding region is denaturated in that particular condition. Two groups of peaks appear at different urea concentrations: one around 3 M and another one at ~6 M, suggesting there are at least two different regions of the protein, whose unfolding is independent and exhibits moderate cooperativity (Figure 2). The maximum number of resonance lines appeared at 9 M urea (Figure 2 – 9 M urea), so this condition was chosen for the assignment of the NMR spectrum.

The DGRASP resonance assignments in 9M urea were based on the two pairs of ^1^H-^15^N-^13^C 3D experiments for sequential linking CBCA(CO)NH + CBCANH and HN(CA)CO + HNCO, in addition to HSQC-TOCSY, HSQC-NOESY-HSQC, (H)CC(CO)NH and HBHA(CBCACO)NH. It was possible to unambiguously assign 96% of the total non-proline HN resonances, together with 95%, 94.5% and 94.4% of the CO, Cα and Cβ, respectively (Figure 3). The assignment transfer to the remaining nine conditions (from 9 to 0 M urea) was done by spectral superposition. Based on this logical approach, all the assignments that could be visually tracked were transferred through the titration spectra. This approach has been successfully used with other non-native molten globule structures [62,63].

**Figure 3:**
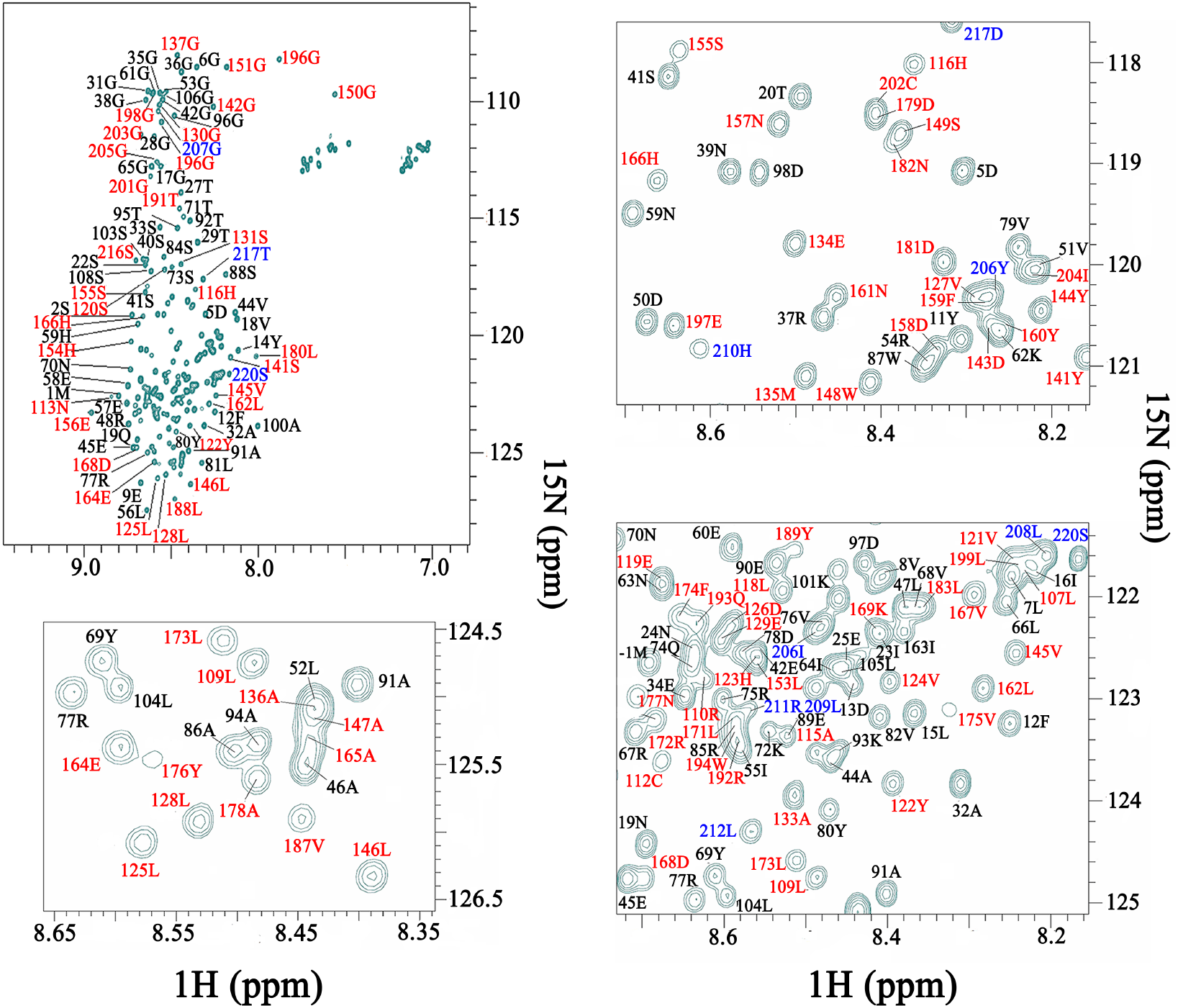
DGRASP resonance assignments: A) ^1^H-^15^N HSQC spectrum of DGRASP in 9 M urea showing the HN resonances that were assigned successfully. Some regions were expanded for better visualization. In black, red and blue are the residues from PDZ1, PDZ2 and SPR, respectively.

Not all peaks were present at the very beginning of the urea titration and only started to appear as the urea concentration was raised (Fig. 2), (a common feature of molten globule states). The gradual appearance of the peaks is attributed to the unfolding of the corresponding region and all the analyses from here on rely on this assumption: if the resonance peak of a non-proline residue X was only observed in the ^1^H-^15^N HSQC at Y M urea, this is the urea concentration where this residue is now assumed to be in a fully unfolded structure. This assumption is valid, since these peaks are all located in a very narrow chemical shift region close to 8 ppm in the proton dimension, typical of unfolded structures and intrinsically disordered proteins [64].

To further explore our data, we monitored every peak that could be unambiguously tracked during the titration, and the urea concentration where each peak appeared during the titration was plotted as a function of the amino acid sequence, giving the local unfolding plot (Figure 4A). Figure 4A clearly shows the differences in the unfolding patterns of the two PDZs in solution, in contrast to the available crystal structures, which suggest that they are structurally identical to each other [12,24]. In general, ~60% of the PDZ1 resonances appear at urea concentrations below 2 M, and the median urea concentration for the unfolding of each amino acid is 2 M (Figure 4A). The most stable part of this subdomain seems to be the region between the amino acids 70 and 95 with urea stability of ca. 4 M. PDZ1 unfolding is therefore weakly cooperative and at least two main unfolding transitions are expected.

**Figure 4.**
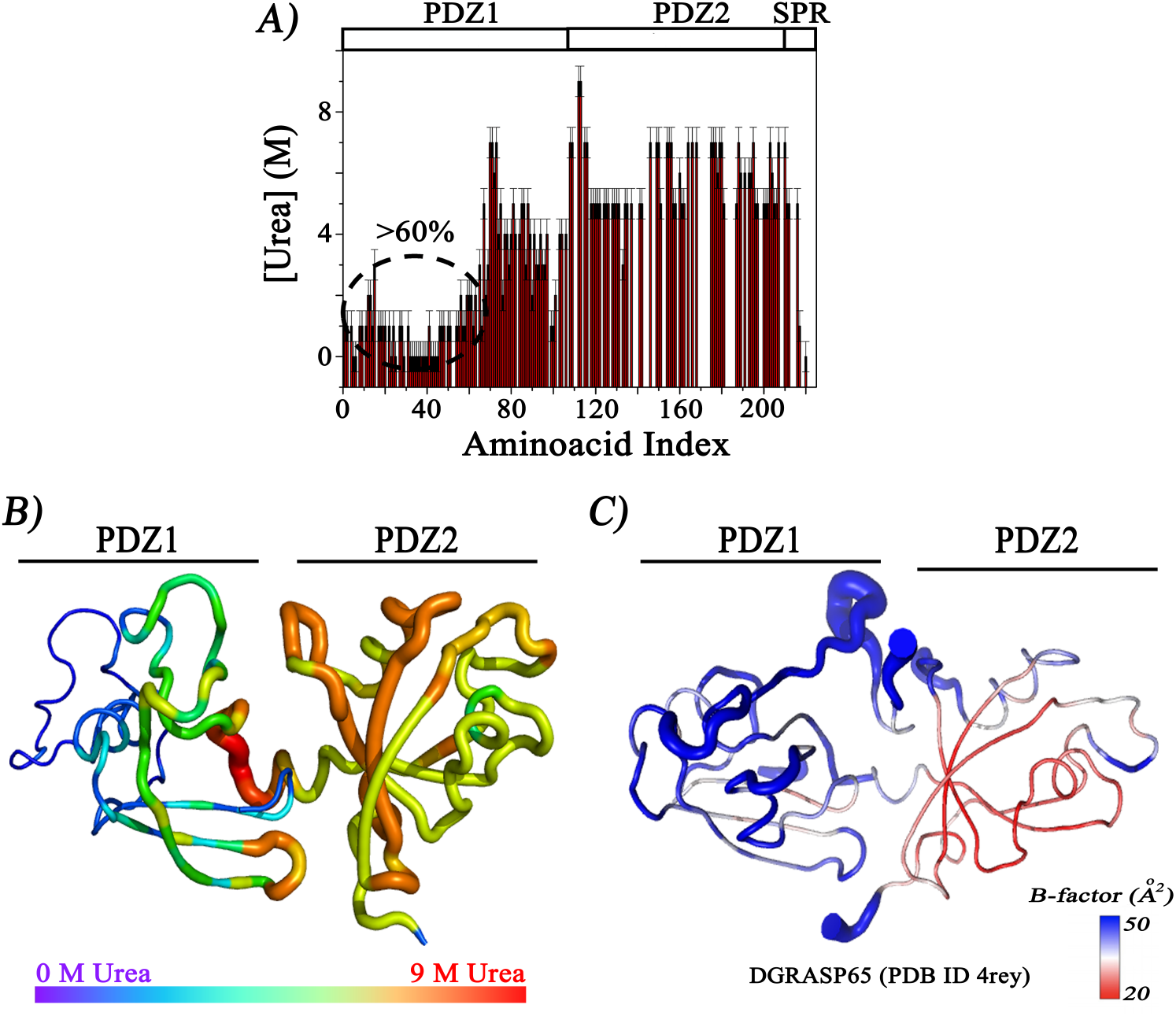
A) Chemical denaturation at residue-specific level. The graphic shows, for each assigned amino acid in the primary sequence, the urea concentration where the resonance line was visible in the ^1^H-^15^N HSQC. The low-stability region inside the PDZ1 was highlighted in the graphic. B) DGRASP molecular model (build on Swiss model [67] and based on the structure of DGRASP65 – PDB I.D. 4rey) highlighted with a gradient of colors according to the urea stability presented in Figure 4A. We assume that the urea concentration necessary to unfold follows a smooth function of variation, so residues without an unambiguous observable urea concentration of unfolding, were given a number based on the simple average of the adjacent amino acids. C) Crystallographic structure of human DGRASP65 showing the experimental B-factors in a gradient of colors. The figures were built in Pymol [68].

Eukaryote PDZs are formed by a β_1_β_2_β_3_α_1_β_4_β_5_α_2_β_6_ secondary structure arrangement [65], but GRASP PDZs have a circular permutation where the first two β-strands are located at the end of the sequence (β_3_α_1_β_4_β_5_α_2_β_6_β_1_β_2_) [12,24]. The PDZ binding groove is therefore formed by the final, rather than the second, β-strand within the fold [24]. PDZ binding grooves are structured into a short α-helix (α_2_) coupled with the abovementioned β_2_ and our NMR data therefore suggest that the PDZ1 binding pocket is formed by a stable β-strand and a more flexible and unstable α-helix (Figure 4B). Since the PDZ interaction mechanism involves the formation of a β-sheet between the β_2_ and the disordered C-terminus of the protein partner [66], a more flexible α-helix might be used to control the overall accessibility to this domain.

The opposite behavior was observed for PDZ2, where the majority of the resonance peaks appeared at 5-7 M urea, in a much sharper transition (Figure 4A-B). The average urea concentration for PDZ2 unfolding is 5.7 M. It is clear that the GRASP domain is formed by the much more stable PDZ2 and a PDZ1 with at least two regions of different stability, where the more unstable one is predominant (>60%) (Figure 4B).

The existing crystal structure of the human GRASP65 GRASP domain (DGRASP65) indicates that PDZ1 is more flexible than PDZ2 based on the B-factor values (Figure 4C). Instability during chemical unfolding may be closely related to structural flexibility, especially in the region close to and surrounding the binding pocket of PDZ1.

### Dissecting the GRASP domain

Using the domain predictor Pfam (http://pfam.xfam.org, accessed in 2017), the disorder prediction published previously [28] and the information from our own solution NMR experiments, we designed constructs for each individual PDZ domain. Residues 1-115 were assigned to “PDZ1” and 116-220 to “PDZ2” and the two constructs were expressed and purified by following the same protocol used for the full-length DGRASP.

SRCD data indicate that the PDZs have different secondary structure content (Figure 5A, Table 1). In solution, PDZ1 possesses lower helical content (approximately three times less than PDZ2) and more strand structures, in contrast to predictions based on the crystal structures, in which similar secondary structure content was observed [24,25]. It has previously been shown that GRASP structure is sensitive to changes in the physicochemical parameters of the medium, in particular the variation in the dielectric constant. [36]. High PEG concentrations have also been shown to induce disorder-to-order transitions in GRASP, so we could expect that more disordered regions might become more ordered in conditions used for crystallization [36].

**Table 1:** SRCD spectral deconvolution performed using the software CDSSTR and an appropriate database in the online server Dichroweb [69].

|  | Helix | Strand | Turns | Disord. | NRMSD |
| --- | --- | --- | --- | --- | --- |
| **PDZ1** | 5% | 37% | 22% | 34% | 0.040 |
| **PDZ2** | 18% | 30% | 22% | 30% | 0.027 |

**Figure 5:**
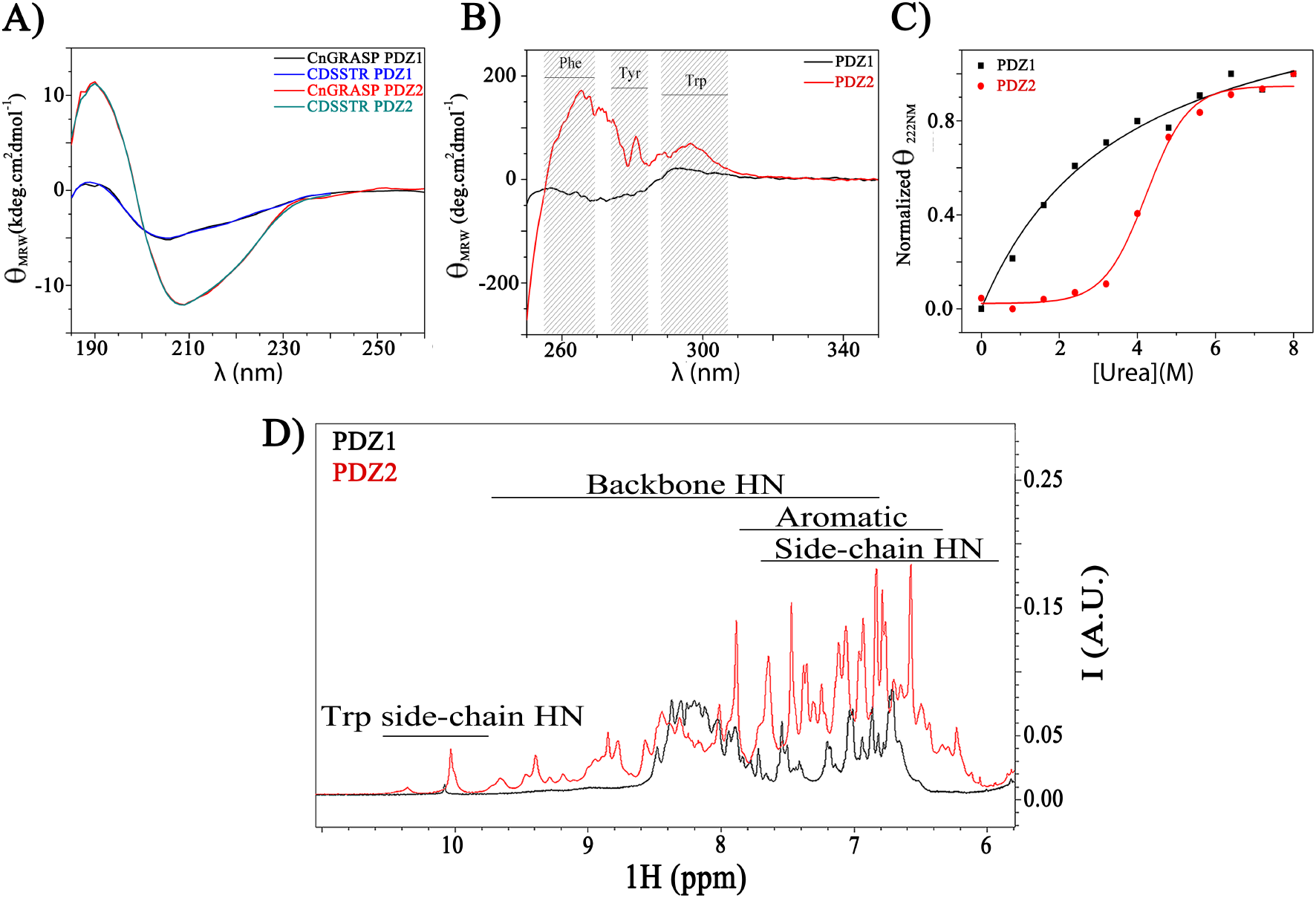
SRCD and ^1^H NMR data for the separate PDZs of CnGRASP. A) SRCD data of PDZ1 (black) and PDZ2 (red) together with the deconvolution fit (blue and green, respectively) obtained using the software CDSSTR and an appropriate database in the online server Dichroweb [70]. B) Near-UV CD of PDZ1 (black) and PDZ2 (red) with the regions of Phe, Tyr and Trp resonances highlighted in grey. C) Urea titration monitored by the intensity at 222 nm using far-UV CD of PDZ1 (black) and PDZ2 (red) as a function of the urea concentration. The PDZ1 and PDZ2 data sets were fitted with a hyperbolic and a sigmoidal function, respectively. D) ^1^H NMR of PDZ1 (black) and PDZ2 (red) with some resonance regions highlighted in the figure for illustration.

Changes in protein tertiary structure may be monitored using near-UV (250-350 nm) CD spectra [71]. If a protein has no well-defined 3D structure (e.g. IDPs or “molten-globule” structures), the signals in the near-UV region will be close to zero [72]. On the other hand, the presence of significant near-UV signals is a good indication that the protein is folded into a well-defined tertiary structure [71].

PDZ1 (PDZ2) has 4 (6) tyrosines, 4 (3) phenylalanines, and 1 (3) tryptophans. Therefore, both PDZs have comparable numbers of aromatic amino acids that are reasonably distributed along the primary structure and that can be used as good probes in near-UV CD experiments. The PDZ2 near-UV CD spectrum has multiple peaks in the region analyzed, suggesting that it has a more well-defined tertiary structure (Figure 5B). The same pattern is not observed for PDZ1, where only a bump is present in the tryptophan region, probably due to its single Trp being in a more collapsed region. The near-UV CD spectra therefore suggest that the isolated PDZ2 has a more well-defined tertiary structure than PDZ1.

Our data so far have revealed some important differences between PDZ1 and PDZ2 that could not be anticipated from the existing crystal structures. Although DGRASP unfolds in a non-cooperative (linear) manner during urea titration (Figure 1), individual PDZs show distinct unfolding transitions when compared to DGRASP and to each other (Figure 5C). PDZ1 starts to unfold very early during the titration (~1 M urea) and, at ~ 2-3 M urea, around 60% of the total population is already in the unfolded state (Figure 5C). This CD data is consistent with the NMR data, in which a significant part of the structure appears to be unfolded at the beginning of the urea titration. It is important to emphasize that CD and NMR provide complementary information: NMR allows us to analyse local denaturation, whereas, in the CD experiment, we observe global structural differences between the folded and the unfolded states. Nevertheless, both NMR and CD data agree that PDZ1 has a less stable structure and with a weakly cooperative unfolding process. In contrast, PDZ2 shows a more strongly cooperative transition to the unfolded state with a urea concentration at half transition of 4.2 M, suggesting that, even though the PDZs have comparable amounts of regular secondary structure and disordered regions, PDZ2 has a more well-defined tertiary structure (consistent with its near-UV CD).

^1^H NMR spectra of proteins are rarely informative because spectral crowding hinders assignment. However, by focusing first on the amide resonance regions, it is possible to see differences between the ^1^H spectra of the individual PDZs (Figure 5D). PDZ2 has multiple peaks spread over the region 6-10.5 ppm, with most of them being very sharp, suggesting that PDZ2 behaves like a regular, ordered protein. A remarkably different behavior is observed for PDZ1 (Figure 5D). The resonances of the amide protons are distributed over a narrower region (~2 ppm) and the line superposition is much more severe. Even though PDZ1 has a poorer dispersion than PDZ2, this dispersion is still greater than those observed in extended intrinsically disordered proteins [73]. Of course, this was expected since neither CnGRASP nor its GRASP domain (DGRASP) is an extended IDP.

The NMR spectra suggest that DGRASP unfolds non-cooperatively and that PDZ2 is more stable, with PDZ1 having at least two regions with different stabilities. Urea titration and NMR experiments with the isolated PDZs validated the initial observations. PDZ1 shows the same pattern observed for DGRASP with fewer peaks at the very beginning of the titration but with a rapid appearance of the remaining lines as the urea concentration is increased (Figure 6A). Most of the peaks missing in the spectrum of the native structure, appeared at urea concentrations around 1-2 M, the same pattern observed in DGRASP (Figure 3B). This unfolding transition is associated with the part of the domain predicted to be the binding site for molecular partners, and it is interesting to note that its unfolding appears to be unaffected by the presence or absence of PDZ2. The remaining lines appeared around 3-4 M urea, suggesting that the most stable region of PDZ1 in the DGRASP construct decreases its stability in the absence of PDZ2 (from ~5 to 3-4 M urea). Interestingly, if we superimpose the PDZ1 and DGRASP spectra acquired in the absence of urea, it is clear that all resonances present for the DGRASP at the beginning of the titration correspond to PDZ1 residues (Figure S2). As discussed before, this suggests these regions are within disordered/flexible structures, and supports the transfer of the assignments from the 9 M urea condition.

**Figure 6:**
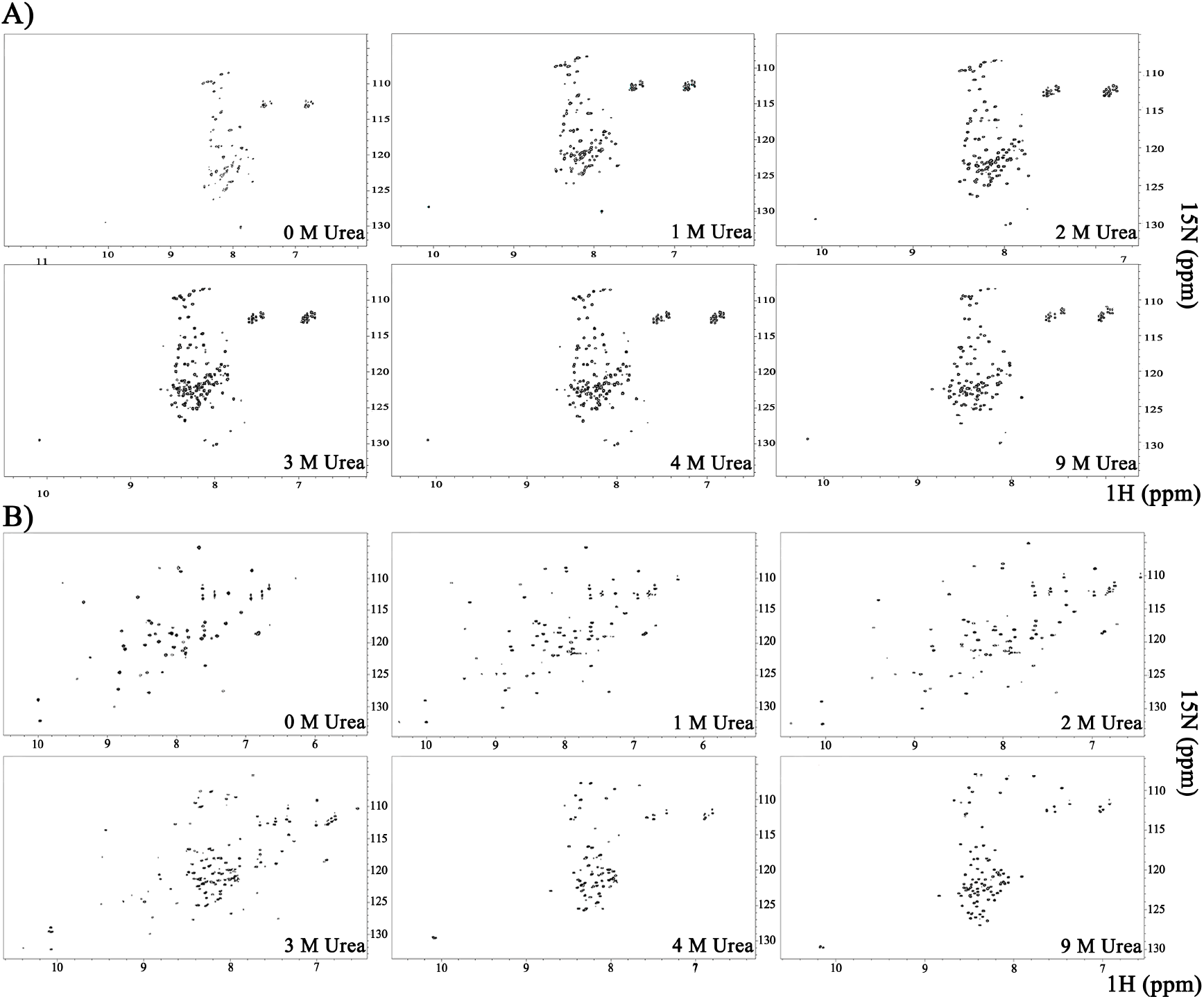
Urea titration experiments monitored by ^1^H-^15^N HSQC in 950 MHz at 20°C. A) Isolated PDZ1. B) Isolated PDZ2.

For PDZ2, the profile is consistent with a regular well-folded protein. The spectrum of the native structure has a reasonable spread of peaks in the proton dimension and, when the urea concentration reaches a specific value, the lines move to a very narrow region close to the ^1^H chemical shift of 8 ppm (Figure 6B). The transition between these two states is therefore highly cooperative and occurs at a urea concentration of around 3-4 M. The cooperative pattern is the same observed previously for the DGRASP but the overall stability is significantly reduced (by at least 1 M urea). Surprisingly, PDZ2 is therefore even more stable when it is connected to PDZ1 in tandem.

### The long-range coupled mechanism of action

Our NMR and SRCD data show that the PDZ domains of GRASPs have different structural stabilities and dynamics, but it is not clear how these properties are exploited to enable the protein to bind multiple different partners. In the crystal structure of the DGRASP55 and JAM-C complex (PDB ID 5GMI), the C-terminus of JAM-C is buried in the PDZ1 binding pocket, however the remainder of JAM-C forms multiple additional interactions with both PDZ1 and PDZ2 [74]. In contrast corresponding crystal structures of DGRASP55 in complex with Golgin45 and the JAM-B C-terminus (PDB IDs 5H3J and 5GMJ) show different molecular contacts with the PDZs [74,75].

To understand the cellular role of PDZ2, we performed MD simulations to sample the conformational flexibility of DGRASP55 in the presence (holo) and absence (apo) of a protein partner (the JAM-C C-terminus peptide) (Figure 7A). Using Boltzmann statistics, for each increase of 1 kcal/mol in free energy at 310 K, a decrease of 80% probability for the observation of the higher energy states is expected. A free energy barrier of approximately 3 kcal/mol would represent an upper limit for the observable energy states at 310 K (with about 1% probability of occupancy). The conformational free energy profile of DGRASP55 in the apo form at physiological temperature (310 K) is large, however when the protein partner is bound (holo) the profile is greatly reduced (Figure 7B). The data suggest that the protein partner significantly reduces the conformational space that DGRASPs can access by stabilizing one particular conformation.

**Figure 7:**
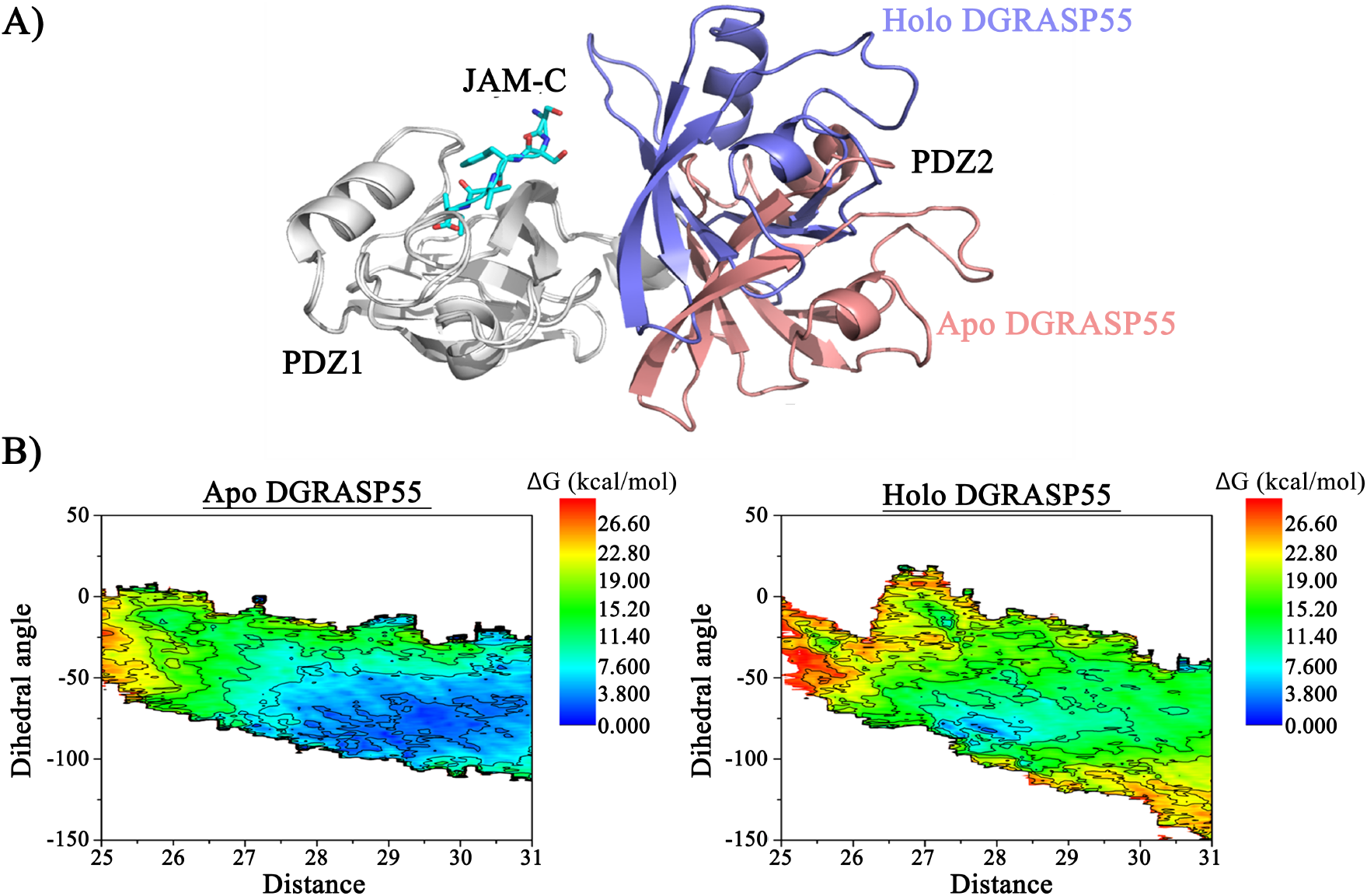
(A) Superposition of crystallographic models of DGRASP55 in the presence (holo-DGRASP55) and in the absence (apo-DGRASP55) of JAM-C C-terminus peptide. The structures were aligned using only PDZ1 (represented in white). Holo-DGRASP55 exhibited a closed conformation characterized by a 33° rotation angle of the PDZ2 towards the PDZ1 in comparison with the apo structure. Apo and holo DGRASP55 PDZ2 are displayed in pink and blue, respectively. The peptide is represented as sticks and colored in cyan. (B) Conformational free energy profile of apo and holo DRASP55. The large accessible area of the apo free energy profile implies a greater conformational flexibility.

Previous crystallographic data have suggested that DGRASP55 transitions from an open to a closed conformation on binding JAM-C and JAM-B [74], however, this closure is not observed when DGRASP55 binds the C-terminus of Golgin45 [75], suggesting that the conformational change, if it really occurs in solution, may be partner dependent. The magnitude of the conformational change (a relative rotation of the PDZs) is sampled in the MD simulations to generate a free energy profile at physiological temperature (Figure 8). The MD simulations were performed using the crystal structures of holo DGRASP55+JAM-C, while the apo free energy landscapes were simulated from the holo structures by removing the peptide (Figure 7B). Under these conditions, the crystal structure conformation observed in the apo forms (PDB ID 4KFW and the homologue 3RLE) was accessible, as were the conformations observed in the complexes with Golgin45 (PDB ID 5H3J) and JAM-B (PDB ID 5GMJ) (Figure 8). At physiological temperature the conformational flexibility of DGRASPs allows large movements in the relative orientations of the two domains, allowing different conformations to be adopted for different binding partners, thereby creating numerous potential secondary interactions via PDZ2. This effect, coupled to the promiscuity of the PDZ1 domain, enables DGRASPs to interact with multiple different protein partners, allowing it to fulfill a key role in cellular signaling.

**Figure 8:**
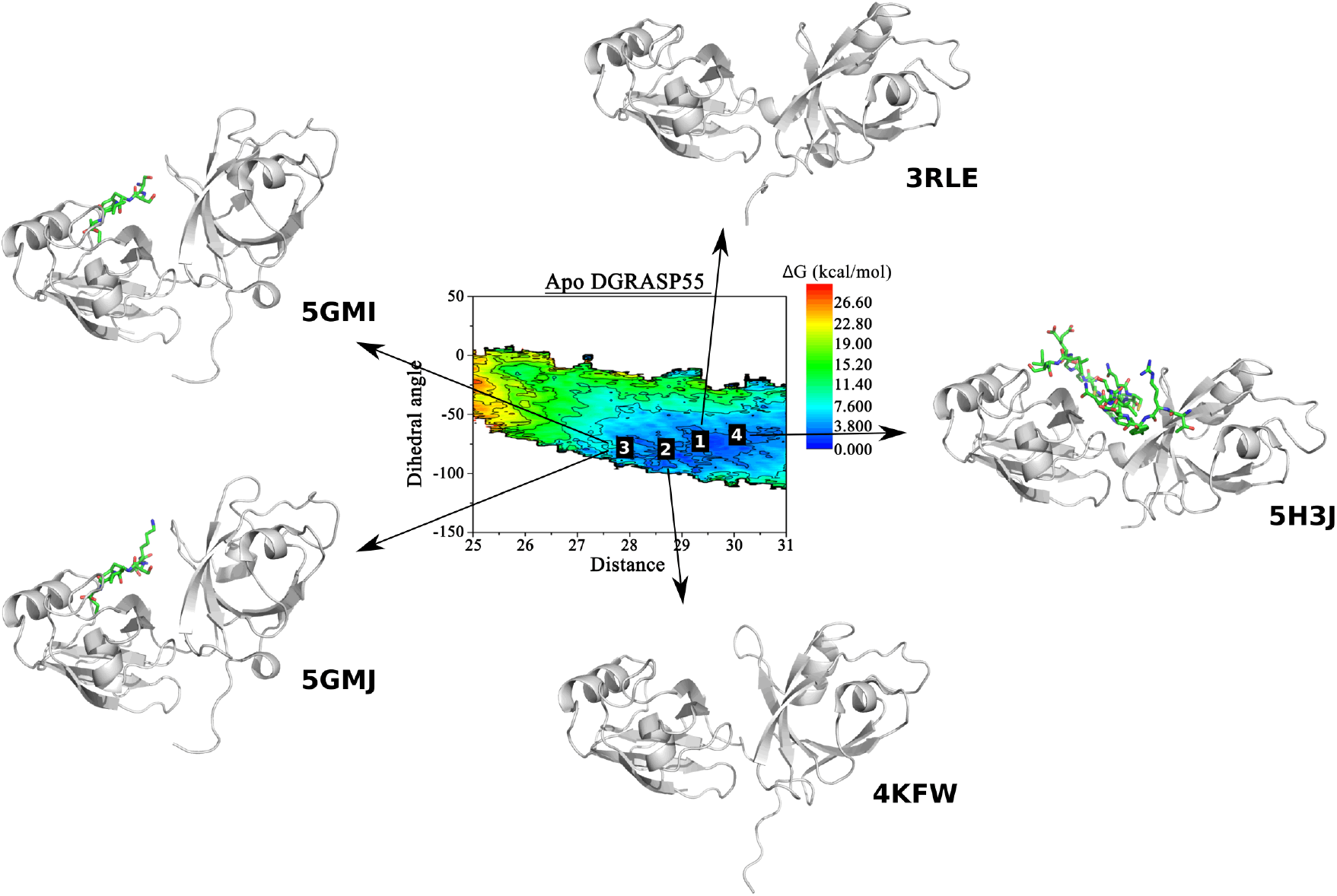
Available DGRAS55 crystallographic models and their coordinates in the free energy map: all the conformations observed in the experimental models are located at the accessible free energy area at physiological temperature for apo-DGRASP55.

## Conclusion

GRASPs are formed by two asymmetric PDZs domains: PDZ1 is more unstable and flexible while PDZ2 is much more stable and structurally more well behaved. Many of the unstable regions found in PDZ1 are located in the binding pocket, suggesting that structural promiscuity inside this domain allows the binding of multiple protein partners. In contrast, PDZ2 is more structured, and this correlates with the smaller number of associated proteins capable of anchoring with GRASPs using this domain. Our findings challenge the conclusions drawn from existing crystal structures, which suggest that both PDZs have a similar structure, even though their function is remarkably different. We have shown that DGRASPs exhibit a large conformational flexibility at physiological temperature, which reduces after binding to a specific protein partner. The coupled, coordinated movement between the two PDZs and the structural promiscuity of PDZ1 appears to enable GRASPs to interact with multiple partners, allowing it to act as a multitasking protein in the maintenance and remodeling of the Golgi body, as well as playing a key role in both conventional and unconventional secretory pathways.

## Acknowledgments

The authors thank the Brazilian agencies Conselho Nacional de Desenvolvimento Científico e Tecnológico (CNPq) and Fundação de Amparo à Pesquisa do Estado de São Paulo (FAPESP) for the financial support through grants no. 2015/50366-7, 2012/20367-3, and 308380/2013-4. LFSM acknowledges FAPESP for the post-doctoral grants (Grant No. 2017/24669-8) and for the FAPESP Research Internships Abroad grant (Grant. No. 2016/09676-5). MRBB acknowledges FAPESP for the post-doctoral grant No. 2016/16328-3. AW acknowledges the UK BBSRC (research grant number BB/N006011/1). We are grateful for the award of SRCD experimental time (SM16289) on the B23 Beamline at Diamond Light Source (UK) under the Membrane Protein Laboratory, funded by the Wellcome Trust grant No. 20289/Z16/Z and we acknowledge the support of Dr Giuliano Siligardi, Dr Rohanah Hussein and Dr Charlotte Hughes. The Bruker console and probe upgrade to the 950 MHz NMR spectrometer was funded in Oxford with support from the Wellcome Institutional Strategy Support Fund, the John Fell Fund and the EPA Cephalosporin Fund.

## Author contributions

LFSM, AJCF, AW and CR designed research. LFSM collected all the experimental data. MRBB performed all the molecular dynamic simulations. LFSM, CR, AW, PJJ and AJCF performed the analyses. AJCF and AW conceived and coordinated the project. All authors contributed to the writing and reviewing of the results. All authors also approved the final version of the manuscript.

## Supplementary Information

**Figure S1:**
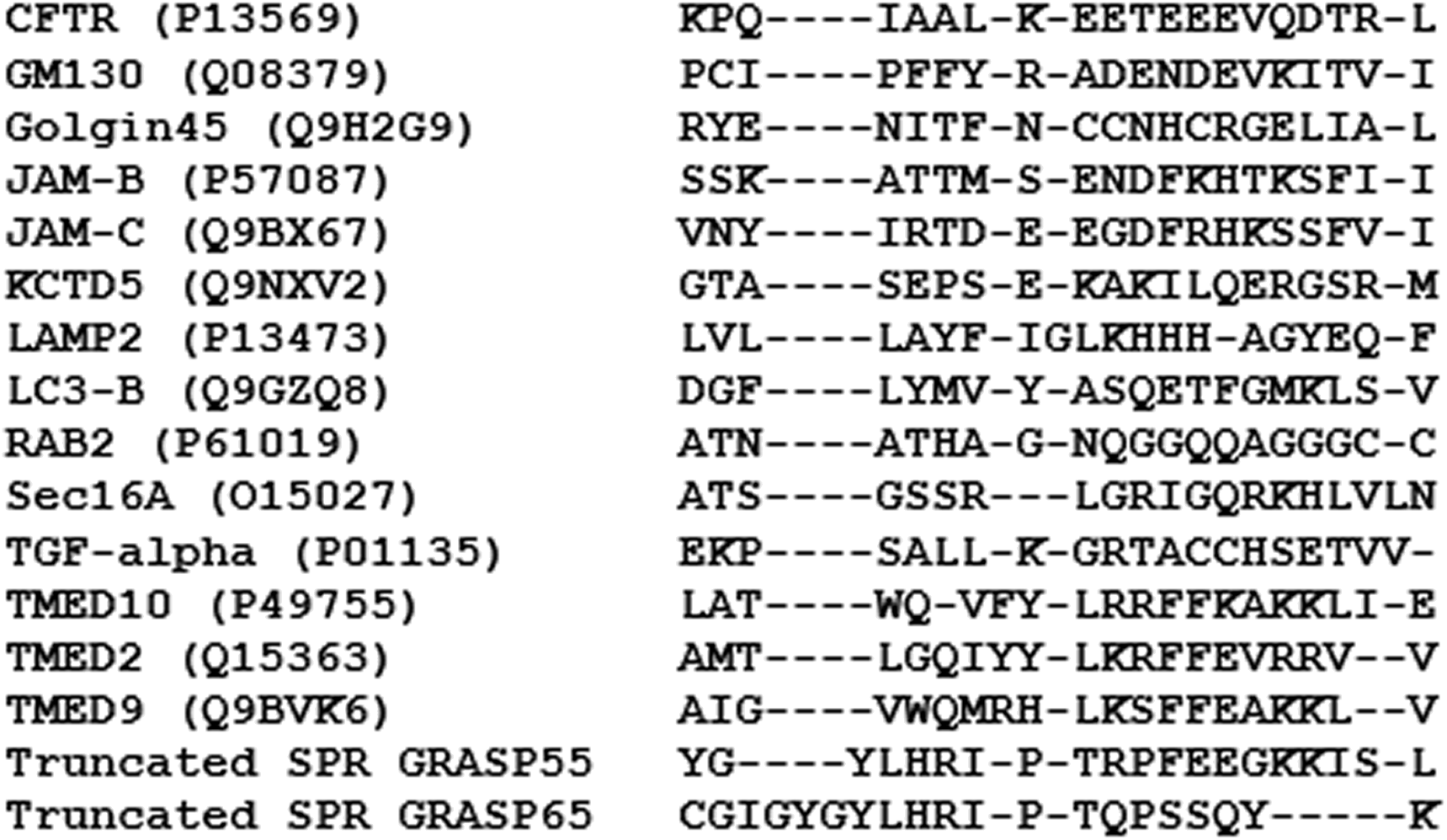
The C-terminus of different GRASP55/65 proteins partners were aligned using T-Coffee (https://www.ebi.ac.uk/Tools/msa/tcoffee/ accessed in 2018).

**Figure S2:**
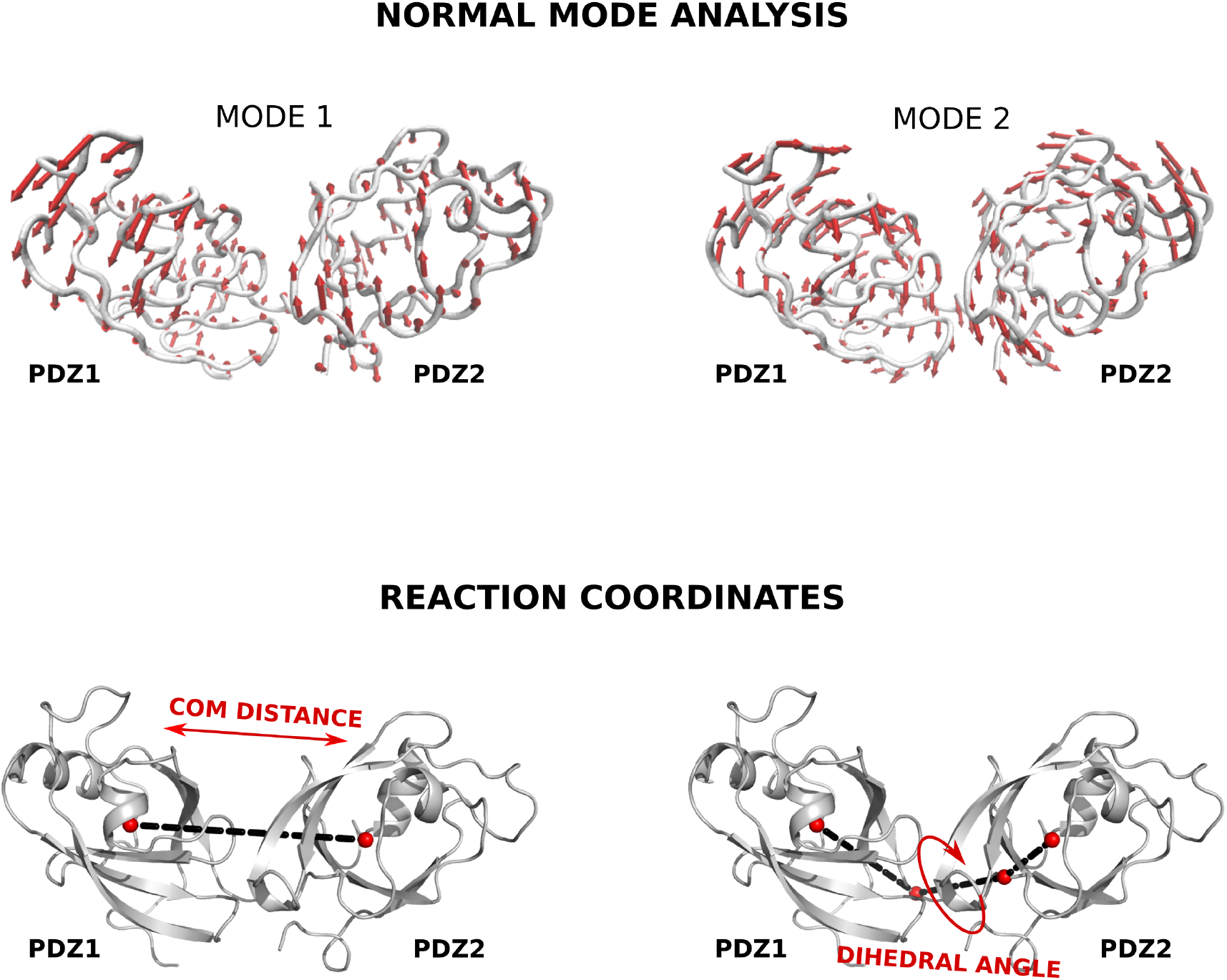
The reactions coordinates used in ABF simulations were chosen based on normal mode analysis (NMA). Briefly, normal modes describe the protein’s natural motions, called normal modes and their associated frequencies. Biologically relevant modes, i.e, modes with low frequency are associated with large-scale motions. Additionally, if there are two known conformations of a protein, the normal modes that most contribute to the conformational change can be used to understand the pathway and to describe the conformational transitions in only a few degrees of freedom. To characterize GRASP collective motions, we performed ANM (Anisotropic Normal Mode) analysis using ProDy [Ref S1]. Visualization of the motion along the two modes with lower frequencies reveals opening/closing and twisting movements. Based on this analysis, the first reaction coordinates coordinate was defined as distance between the center of mass of the two PDZs to capture the opening/closing movements and the second reaction coordinate used in ABF simulations was set as the dihedral angle formed by the center of mass of the PDZs and the helix that connects the two domains to reproduce the twisting motions.

